# DISSECT: A new tool for analyzing extremely large genomic datasets

**DOI:** 10.1101/020453

**Authors:** Oriol Canela-Xandri, Andy Law, Alan Gray, John A. Woolliams, Albert Tenesa

## Abstract

Computational tools are quickly becoming the main bottleneck to analyze large-scale genomic and genetic data. This big-data problem, affecting a wide range of fields, is becoming more acute with the fast increase of data available. To address it, we developed DISSECT, a new, easy to use, and freely available software able to exploit the parallel computer architectures of supercomputers to perform a wide range of genomic and epidemiologic analyses which currently can only be carried out on reduced sample sizes or in restricted conditions. We showcased our new tool by addressing the challenge of predicting phenotypes from genotype data in human populations using Mixed Linear Model analysis. We analyzed simulated traits from half a million individuals genotyped for 590,004 SNPs using the combined computational power of 8,400 processor cores. We found that prediction accuracies in excess of 80% of the theoretical maximum could be achieved with large numbers of training individuals.

## Introduction

The astonishing rate at which genomic and genetic data is generated is rapidly propelling genomics and genetics research into the realm of big data^1^. This great opportunity is also becoming a big challenge because success in extracting useful information will depend on our ability to properly analyze extremely large datasets. The problems associated with big data become critical when, for instance, fitting Mixed-Linear Models (MLMs) and performing Principal Component Analyses (PCA)^2–9^. These analyses are used in a wide range of fields ranging from predictive medicine and epidemiology, to animal and plant breeding, or pharmacogenomics. However, when they are applied to large datasets, one needs to apply workarounds such as performing approximations^3,8^, restricting the applicability to particular cases^9^, and often, even the workarounds need at least one highly computationally demanding step^5^. Furthermore, these workarounds are not scalable. That is, they cannot accommodate increasing compute workloads and volumes of data because they are limited by the memory and computational power available within a single computer. As has happened in other fields^1^, to overcome these limitations the next step must be to move to software capable of combining the computational power of thousands of processor cores distributed across the compute nodes of large supercomputers.

To fill this gap, we developed DISSECT (http://www.dissect.ed.ac.uk/), a highly scalable, easy-to-use and freely available tool able to perform a large variety of genomic analyses with huge numbers of individuals using supercomputers. We showcase our tool by addressing the challenge of predicting phenotypes from genotype data in unrelated human populations. Phenotypic prediction is of central interest to many disciplines and is one of the driving forces behind large-scale genotyping and sequencing projects in a wide range of species^10–14^. Despite considerable efforts, predicting complex traits in unrelated humans has been an elusive goal^12,15^. Accurate prediction of complex traits is expected to be strongly dependent on the availability of sufficiently large datasets^11,15,16^ and the capacity to analyze them together, therefore this being a good challenge to show DISSECT’s capabilities. With this in mind, we simulated a cohort of up to half a million individuals and used DISSECT and the aggregated power of up to 8,400 processor cores to analyze it. We showed that MLMs could predict quantitative traits with increasing accuracy as the sample size of the training cohort increased, and achieved over 80% of the theoretical maximum accuracy when the training cohort had 470,000 individuals. Interestingly, our results also showed that the noise introduced by increasing SNP density has a detrimental effect on the prediction accuracy thus indicating that this increase may not always be desirable.

## Results

### Overview of DISSECT

DISSECT can take advantage of the aggregate power of thousands of processor cores available in supercomputers to perform a wide range of genomic analyses with very large sample sizes. It does that by distributing both data and computations over multiple networked compute nodes that share the computational task, each node having access to only a small portion of the data. Therefore, this computational approach is necessarily more involved than parallelization for desktops, workstations, or single compute nodes on a cluster (in the following text these will be referred to as a single compute node). In addition, the distribution of workload introduces a relative loss of computational power due to the need for communication between compute nodes, which is limited by the speed of the network connection. However, its broad scalability enables the analysis of datasets of sizes that are well beyond the computing capacity of a single compute node, and importantly it does it without the need of performing any mathematical approximation. DISSECT can also analyze moderately large sample sizes with considerably reduced computational time, or run on a single computer when the sample size and computational requirements of the analyses do not require a supercomputer. DISSECT linear algebra computations are based on optimized versions of the ScaLAPACK^17^ libraries to ensure optimal computational performance.

DISSECT implements several highly computational demanding analyses. Some of the most relevant are: performing PCA for studying population structure in large datasets; fitting univariate MLMs; fitting bivariate MLMs, which greatly increase power to detect pleiotropic loci^18^, but require a computational time that is rougly eight times bigger than fitting univariate MLMs to datasets of the same size; regional MLM fitting for studying the accumulated variance explained by the alleles within genomic regions^19,20^, each region having similar computational cost regardless of the number of SNPs fitted but requiring and independent fit; standard regression models with very large number of fixed effects (i.e. fitting the markers of a whole chromosome as fixed effects when extremely large sample sizes are available). DISSECT also allows other computationally less demanding analyses such as the prediction of individual phenotypes from estimated marker effects (i.e. polygenic scores^21^) or standard GWAS analyses. Furthermore, it also implements optimized routines similar to those found in GEMMA^5^ which allow DISSECT to run much faster analyses with less resources when it is possible. For instance, by diagonalizing the covariance matrix, thus enabling fast MLM fitting.

### Computational performance

We performed MLM and PCA analyses using simulated cohorts (Online Methods) of different sample size (N) (Fig. 1) to show the computational capabilities of DISSECT. We selected these two examples because they are highly computational demanding analyses, requiring a running time of O(N^3^). The analyses were run on the UK National Supercomputing Service (ARCHER), a supercomputer with 4,920 computer nodes containing 9,840 processors with 12 cores each (i.e. a total of 118,080 cores available). DISSECT is able to fit, after eight iterations, a MLM to a sample of 470,000 individuals and 590,004 SNPs in less than four hours using the aggregated power of 8,400 cores and a total of ∼16TB of memory (∼2GB of memory per core). The running time included estimation of the variances using REML^22,23^, best linear predictions of the individual’s genetic values and best linear predictions of SNP effects^24,25^. If we disregard the computational performance loss due to communication between nodes, we can roughly estimate the computational time required by a computer with one core to complete the analysis by multiplying the number of used cores with the computation time (core-hours). In this situation, the MLM fit would need 3.6 years (Fig. 1a). Performing a PCA for 108,000 individuals and same number of SNPs, required ∼2 hours using 1,920 cores. That is, arround ∼4,000 core-hours which would be equivalent to ∼160 days of computation on a single core (Fig. 1b). All these results show both the high computational demands required for performing these analyses together with the ability of DISSECT to perform them.

**Figure 1:**
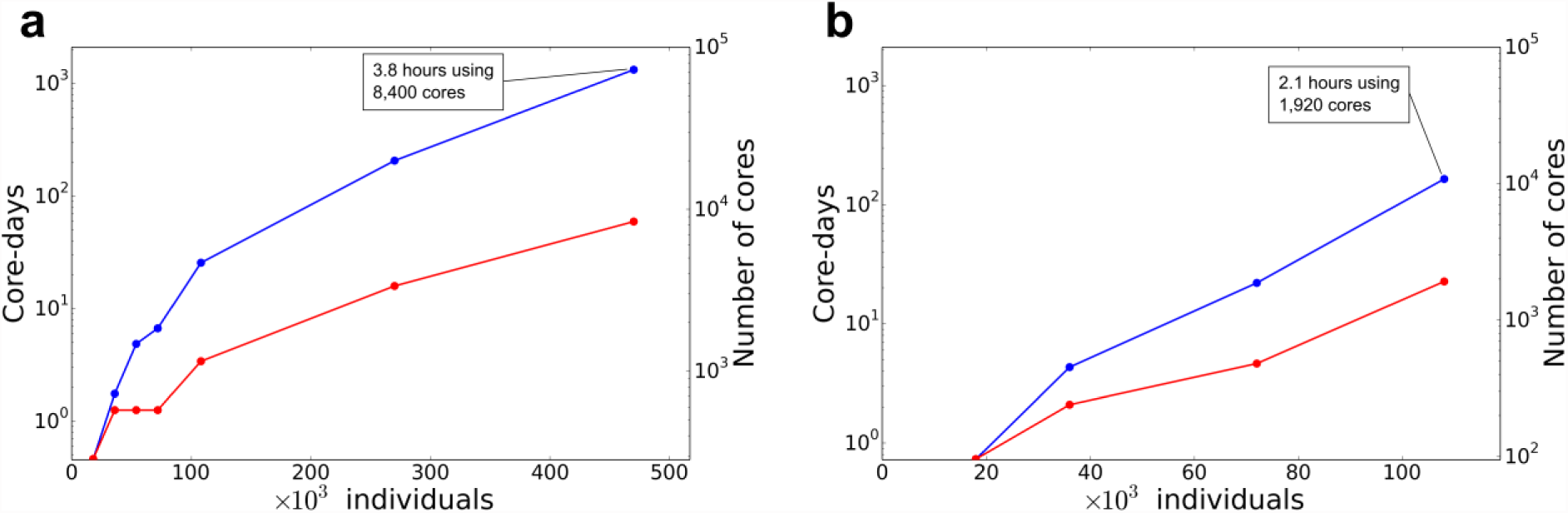
Computational requirements for MLM and PCA. (**a**, **b**) Computational time (blue lines, left axis) and number of processor cores used (red lines, right axis) in log scale for MLM (**a**) and PCA (**b**) analyses as a function of sample size. Core-days is the amount of time in days required to complete the analyses multiplied by the number of cores used. It is a roughly estimate of the computational time a single computer with a single core would require for performing the analyses if DISSECT scaled perfectly (i.e. there was not computational performance penalization due to communication between computer nodes).

### Prediction results with huge sample sizes

We tested the accuracy of phenotypic prediction from genotype data when large numbers of individuals are available. To this end, more than half a million SNP genotypes for over half a million individuals were simulated based on linkage disequilibrium (LD) patterns and allele frequencies of Hapmap CEU population. Then, we simulated several quantitative traits by using both, different heritabilities (h^2^), and numbers of quantitative trait nucleotides (QTNs). In each case, we divided the cohort in two subsets, one for training the models and another for validating the predictions (Online Methods). Predictions were based on the effects of all available SNPs estimated jointly from the MLM fit. As expected, prediction accuracy increased with the heritability of the trait and the size of the training dataset (Fig. 2). The MLM efficiently captured the effects of large numbers of genotyped and ungenotyped QTNs and its performance was unaffected by the number of QTNs affecting the trait (Fig. 2 and Supplementary Fig. 1). Importantly, high accuracies were only achieved when large numbers of individuals were used to train the prediction model. For instance, training the MLM with 470,000 individuals yielded correlations of 0.72, 0.57, and 0.30 for traits with 10,000 QTNs and heritabilities of 0.7, 0.5, and 0.2, respectively. That is, between 86% and 68% of the theoretical maximum, which is the square root of the heritability. Simulated traits determined by 1,000 QTNs gave very similar results to traits determined by 10,000 QTNs (Supplementary Fig. 1). We explored why even when training the models with this extremely large sample sizes, the limit of prediction accuracy was yet not close to the theoretical maximum. Estimation of QTN effects is very accurate (Supplementary Fig. 2), therefore we hypothesised that the loss in accuracy could be due to QTNs not being properly tagged by markers in the array, or due to the the noise introduced by the linkage disequilibrium structure of the genome.

**Figure 2:**
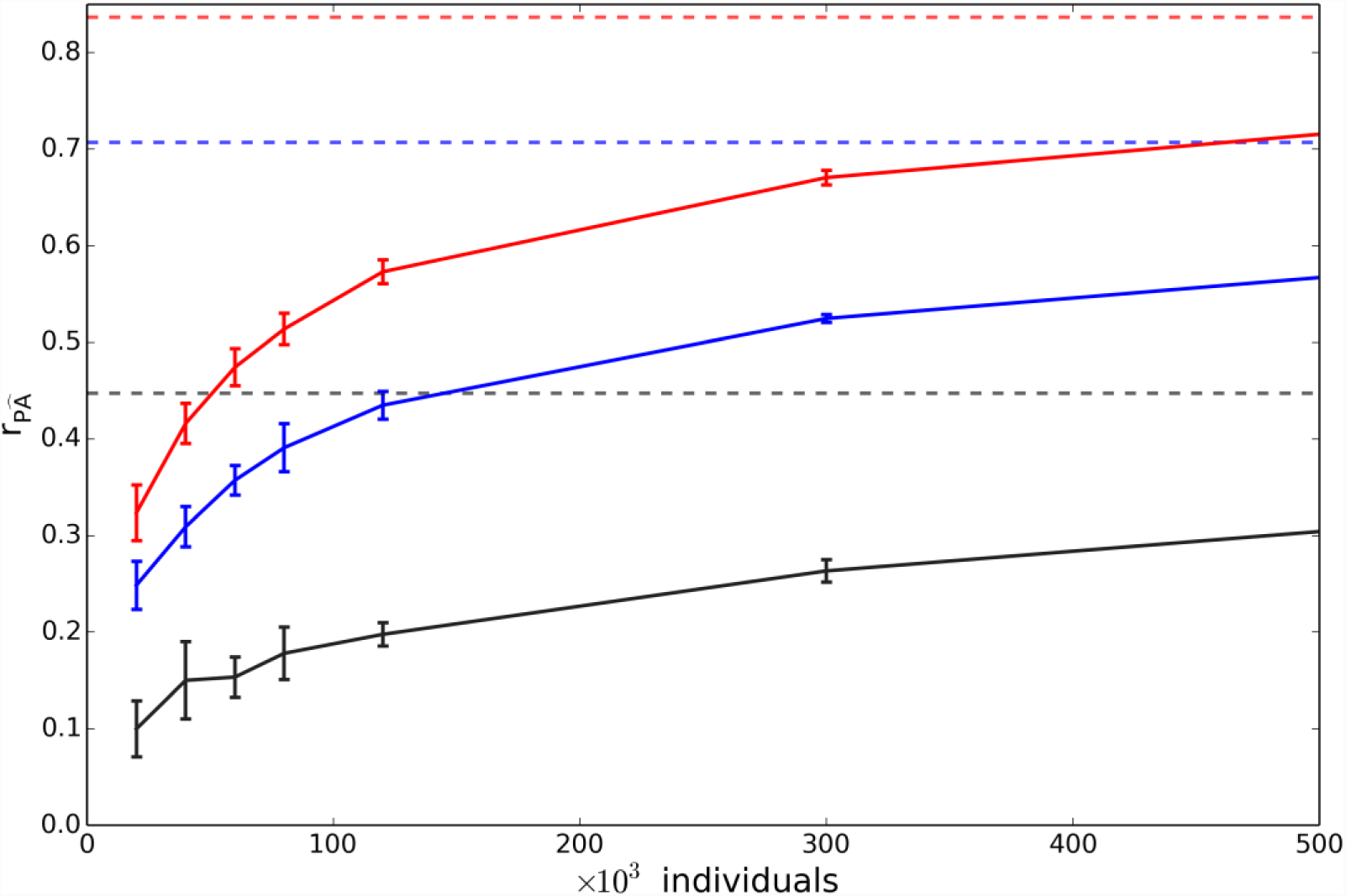
Prediction accuracy of MLM as a function of sample size and heritability. Correlation between true and predicted phenotypes as a function of cohort size for a trait determined by 10,000 QTNs. Black, blue and red curves represent heritabilities of 0.2, 0.5, and 0.7, respectively. Constant dashed lines indicate the theoretical maximum achievable for each heritability. Error bars are two times the standard deviation.

### Prediction accuracy when all QTNs are genotyped

An important question is whether one could reach the theoretical limit of prediction accuracy by genotyping or sequencing all QTNs^26^ whilst being unable to discriminate causal from non-causal variants. We simulated new phenotypes assuming the genotypes for all QTNs were included in the genotyping array. We repeated all our previous analysis and showed that the prediction accuracy for traits with 10,000 QTNs increased only slightly (Fig. 3). Traits with 1,000 QTNs give very similar results (Supplementary Fig. 3). Since this increase was not as high as we expected, it raises serious doubts that genotyping or sequencing the QTNs will improve prediction accuracy if the QTNs effects cannot be disentangled from the effects of other correlated SNPs (Supplementary Fig. 4). These results indicate that the noise introduced by SNPs that are not QTNs significantly reduce the accuracy of prediction, even for very large number of individuals.

**Figure 3:**
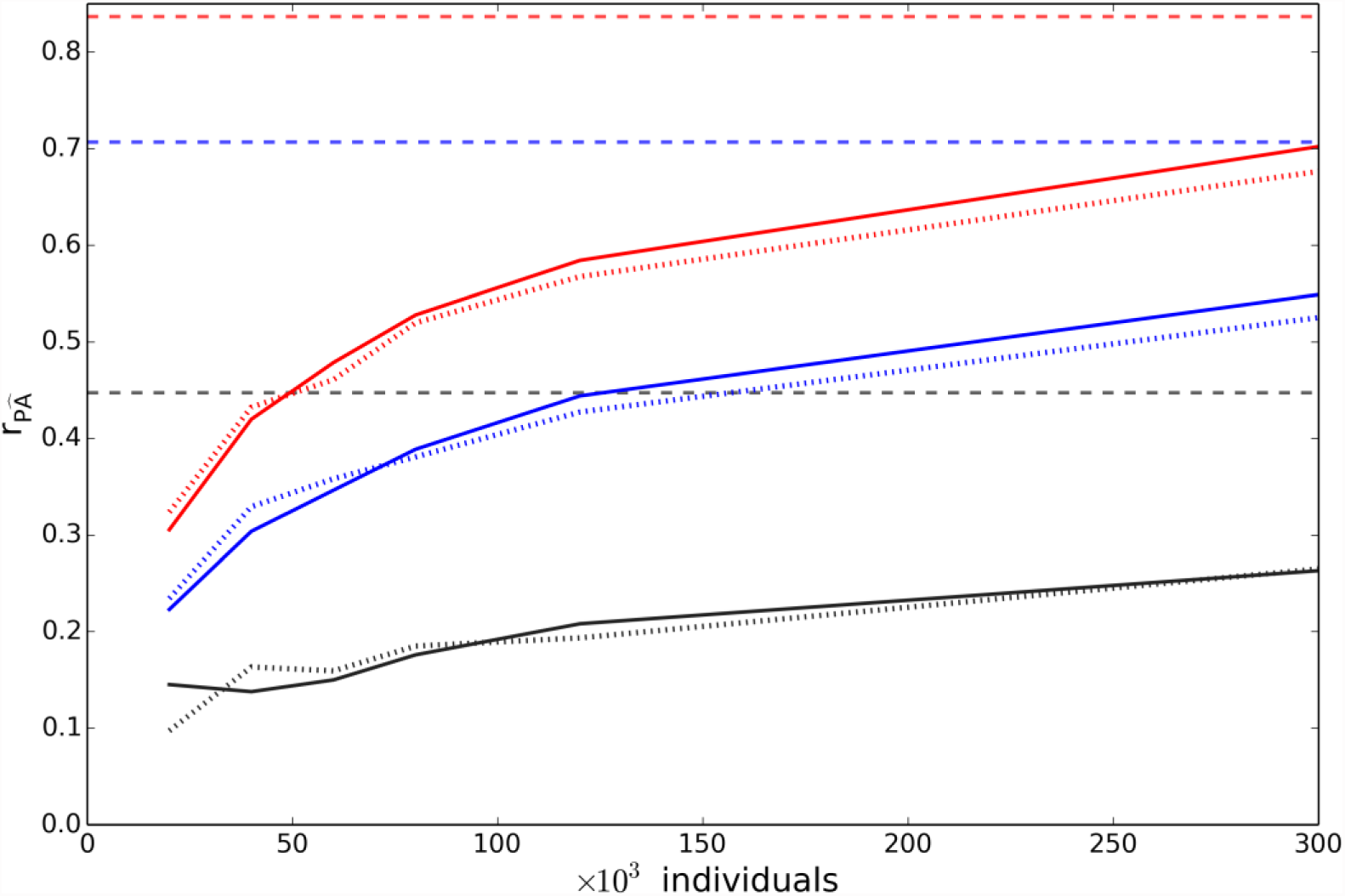
Prediction accuracy when all QTNs were genotyped. Correlation between true and predicted phenotypes as a function of the cohort size when the trait is determined by 10,000 QTNs. Black, blue and red curves represent traits with heritabilities of 0.2, 0.5, and 0.7, respectively. Solid lines are the correlations obtained when all QTNs were genotyped. Dotted lines are the correlations obtained when only ∼20% of QTNs were genotyped. Constant dashed lines indicate the maximum theoretical correlation for each heritability.

## Discussion

We have presented DISSECT, a new tool to perform a wide range of genetic and genomic analyses that overcomes the limitations of single compute nodes, when huge sample sizes are available, without the need of performing approximations or compromises in terms of the model fitted to the data. An ever more pressing need if one considers the release of very large genotyped cohorts like the UK Biobank.

We showcased DISSECT by addressing the timely topic of complex trait phenotypic prediction, which is of central interest to many disciplines. Prediction in unrelated humans has been an elusive goal^12,15^ due to a combination of suboptimal statistical methodology, small training datasets, and lack of computational tools. DISSECT allowed us to fit MLMs to near 500,000 individuals and around 600,000 SNPs reaching prediction accuracies of up to 80% of the theoretical maximum on simulated quantitative traits. We also have shown that the noise introduced by highly correlated SNPs has a strong impact on the accuracy of prediction when using MLMs for prediction, and therefore increasing SNP density could have an adverse effect on the accuracy of prediction even for extremely large sample sizes.

Although we showcased DISSECT by addressing the problem of phenotypic prediction in humans, it can also be used in plant and animal breeding and perform a wide range of commonly used analyses. In addition, DISSECT is under active development and there are several new functionalities planned or in testing stage.

## Methods

### Simulations

We used the HAPGEN 2 software^27^ for simulating half a million individuals -based on linkage disequilibrium (LD) patterns and allele frequencies of 2,543,887 SNPs available in the Hapmap 2 (release 22) CEU population^28^-from which we generated subsets of 20, 40, 60, 80, 120, 300, and 500 thousand individuals. From each subset of data, we used 90% of the individuals for training the models and the rest for validating the predictions. The only exception was the subset including 500,000 individuals, where we used 470,000 individuals for training and 30,000 for validation. We simulated polygenic and highly polygenic quantitative traits that were determined by 1,000 and 10,000 randomly distributed quantitative trait nucleotides (QTNs), respectively. The QTNs were randomly distributed across the genome and their combined effects explained 20, 50 and 70% of the phenotypic variation. That is, we simulated heritabilities (h^2^) of 0.2, 0.5, and 0.7. The QTNs effects were the same for all data subsets. Six replicates were performed for each trait heritability and genetic architecture. Each replica assumed different QTNs with different random effects.

The simulations were performed using DISSECT assuming an additive genetic model:

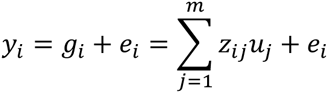

with y_i_ being the quantitative trait of individual i, u_j_ the effect of QTN j drawn from a normal distribution with mean zero and variance one, m the number of assumed QTNs and e_i_ a normal distributed random variable with zero mean and variance 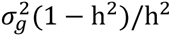 where 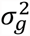 is the variance of g_i_. z_ij_ is the normalized genotype of individual i at QTN j. It is defined as *z*_*ij*_ = (*s*_*ij*_ – μ_*j*_) / σ _*j*_ where s_ij_ is the number of reference alleles at QTN j of individual i, μ_*j*_ = 2*p*_*j*_ and 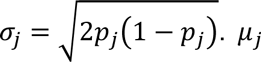 and σ_*j*_ are the mean and the standard deviation of the reference allele among the individuals genotyped, defined as a function of the reference allele frequency (p_j_).

### MLM and Prediction

MLMs analyses were performed using DISSECT. For our first set of analyses we excluded all SNPs not present on the Illumina Human OmniExpress BeadChip (i.e., we analyzed a total of 590,004 SNPs), that is only ∼20% of the QTNs were genotyped. Later, we investigated the effect of having the QTNs in the genotyping array and included the remaining ∼80% of QTNs to the genotyping array.

The model fitted was:

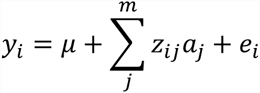

where *μ* is the mean term and e_i_ the residual. z_ij_ is the normalized genotype of individual i at QTN j. The vector of random SNP effects **a** is distributed as 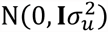. 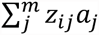 is the total genetic effect for individual i. The phenotypic variance-covariance matrix is 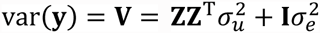. 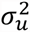 and 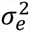 were fitted using the AI REML method^22,23^. SNP effects were estimated using the formula^24^:

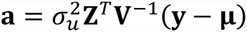

SNP effects were used as an input for DISSECT to predict phenotypes on the validation cohort. DISSECT computes the prediction for individual i as a sum of the product of the SNP effects and the number of reference alleles of the corresponding SNPs:

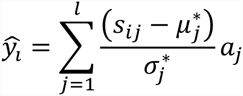

Where s_ij_ is the number of copies of the reference allele at SNP j of individual i, l is the number of SNPs used for the prediction, and a_j_ the effect of SNP j estimated from the MLM analyses. 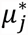 and 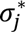 are the mean and the standard deviation of the reference allele in the training population.

## Acknowledgments

This work was mainly supported by the Medical Research Council [grant number MR/K014781/1]; with contributions from Cancer Research UK [C12229/A13154] and The Roslin Institute Strategic Grant funding from the BBSRC. AT also acknowledges funding from the Medical Research Council Human Genetics Unit. We acknowledge Ricardo Pong-Wong for his help. This work used the ARCHER UK National Supercomputing Service (<u>http://www.archer.ac.uk</u>).

## URLs

DISSECT and documentation available at: <u>https://www.dissect.ed.ac.uk</u>

## Contributions

All authors contributed to the conception and design of the study, read and approved the manuscript. OCX wrote the DISSECT software and performed the statistical analyses. OCX and AT wrote the paper.

